# Investigating the performance of Oxford Nanopore long-read sequencing with respect to Illumina microarrays and short-read sequencing

**DOI:** 10.1101/2024.12.19.629409

**Authors:** Renato Santos, Hyunah Lee, Alexander Williams, Anastasia Baffour-Kyei, Claire Troakes, Ammar Al-Chalabi, Gerome Breen, Alfredo Iacoangeli

## Abstract

Oxford Nanopore Technologies (ONT) long-read sequencing (LRS) has emerged as a promising tool for genomic analysis, but comprehensive comparisons with established platforms across diverse datasets remain limited. We present a multi-platform benchmark using 14 human genomes sequenced with ONT LRS, Illumina short-read sequencing (SRS), and Illumina microarrays. Our study evaluates LRS performance for various genetic variants across genomic contexts, while also examining the impact of experimental factors such as multiplexing, depth, and read length. In high-complexity regions, LRS demonstrated competitive yet slightly lower accuracy than SRS for SNV detection (F-measure: 0.954 vs. 0.968), with performance gaps narrowing in low-complexity regions. For indel detection, LRS showed robust performance for small indels (1-5bp) in high-complexity regions (F-measure: 0.869), but accuracy decreased significantly in low-complexity regions and for larger indels. LRS identified 2.86 times more structural variants than SRS, with superior detection of large-scale variations. Sequencing depth strongly influenced variant calling performance across all variant types, while multiplexing effects were minimal after controlling for depth. Our findings provide valuable insights for optimising ONT LRS applications in genomic research and clinical diagnostics.

## Introduction

Since the completion of the Human Genome Project (1), the field of genomics has continually evolved, with newly developed genotyping and sequencing technologies enabling the study of diverse types of genomic variation. The first published human reference genome assembly in April 2003 (2), provided the foundation for various variant calling methods, involving mapping sequenced reads to a reference and identifying nucleotides that differ between the reads and the reference. This advancement has facilitated the study of genomic variations associated with disease and the development of genetic diagnostics in clinical settings (3). Genomic variants can be characterised using several different methods, including DNA microarrays, short-read sequencing, and long-read sequencing, each with its own advantages and disadvantages.

SNP arrays, a type of DNA microarray, have emerged as a powerful and cost-effective tool for genotyping one of the most common types of variation, single nucleotide variants (SNVs) (4). These arrays consist of high-density oligonucleotide probes attached to a solid surface, designed to hybridise with specific DNA sequences containing SNVs of interest. The primary advantages of microarrays include their cost-effectiveness for large-scale studies, high-throughput capabilities, and ability to detect SNVs across the genome with high accuracy (5), leading to their widespread adoption in large-scale population studies and genome-wide association studies (GWAS) (6).

However, microarrays are limited to detecting known variants for which probes have been designed, making them less suitable for discovering novel or rare variants, or structural variation, such as balanced chromosomal rearrangements (5). Despite these limitations, microarrays remain a valuable tool in genomics research, particularly for large-scale genotyping projects where cost and high-resolution throughput are the primary considerations.

Short-read sequencing (SRS) has become a cornerstone technology in genomics research and clinical diagnostics, with widespread adoption due to its high accuracy, throughput, and cost-effectiveness (7). The typical error rates of approximately 0.1% (Q30) (8), allow for confident detection SNVs and small insertions/deletions (indels) in high-confidence regions of the genome, where reads can be accurately mapped (9). Furthermore, recent developments in versatile and scalable bioinformatics analysis pipelines have made it possible to efficiently process and analyse SRS data (10-12). These key advantages have made SRS the go-to technology for both large-scale whole genome studies (13, 14) and clinical applications (15, 16).

Conversely, the relatively short read lengths (typically around 100–250 bp) pose challenges when performing variant calling in repetitive genomic regions, such as segmental duplications (17), transposable elements (TEs) (18), short tandem repeats (STRs) (19), homopolymers (20), and telomeres (9). In these regions, short reads map to the reference ambiguously, or fail to map entirely, leading to incomplete coverage and potential misidentification of variants. Although various computational approaches to improve variant calling in challenging regions have been developed to address these limitations, including methods for local assembly, split-read analysis, and paired-end mapping strategies (21), many dark regions of the genome still remain inaccessible or poorly characterised using SRS alone (22).

In recent years, long-read sequencing (LRS) technologies were developed as powerful complements to short-read sequencing, addressing many of the limitations associated with short reads in repetitive and complex genomic regions. Among these, Oxford Nanopore Technologies (ONT) has gained significant attention due to its unique advantages and potential applications. Introduced commercially in 2014, ONT utilises a fundamentally different approach to sequencing, by passing a single strand of DNA through a protein nanopore embedded in an electrically resistant membrane (23). As the DNA molecule traverses the pore, it causes characteristic disruptions in the ionic current flowing through the pore, which are then measured and translated into base calls (24). This approach allows for the generation of exceptionally long reads, typically ranging from 10 to 100 kb (25), with maximum length reads demonstrated to exceed 2 Mb in length (26). These can span entire repetitive regions, resolve complex structural variants, and improve de novo genome assemblies (22), and allow the detection of epigenetic marks, such as methylation, without the requirement of additional library preparation methods (27).

While initial iterations of ONT LRS had error rates of 38% for R6 nanopore chemistry, based on read-to-reference alignment metrics (28), recent advancements in base-calling algorithms, improved pore chemistry, and sequencing protocols have substantially improved the accuracy of ONT sequencing. Indeed, the latest R10.4 nanopore chemistry has significantly reduced error rates, achieving a modal read accuracy of over 99% (Q20) (29) and an average accuracy of 96.8% (Q15) (30) on read-to-reference alignment. In addition, the development of highly accurate consensus approaches, such as duplex sequencing, has further improved accuracy, albeit at the cost of reduced throughput (31). The ability to generate long reads at increasingly competitive costs and accuracy positions ONT as a valuable tool for comprehensive genome analysis, particularly in regions of low-complexity where short-read technologies fall short, as demonstrated in recent clinical-setting validations (32).

While previous LRS benchmarking efforts have largely relied on well-characterised reference samples from the Genome in a Bottle (GIAB) consortium (32-36), there remains a need for comprehensive comparisons using diverse sample datasets. To address this gap, we present a new benchmark of LRS against other technologies using a cohort of 14 in-house sequenced genomes, while also thoroughly characterising this dataset. Our design utilises a multi-platform strategy, combining data from Illumina microarrays, Illumina SRS, and ONT LRS thereby allowing for a thorough evaluation of ONT’s capabilities in calling SNVs, indels, and structural variants (SVs). Thus, by comparing LRS results to those obtained from well-established technologies, we can gain insights into the strengths and limitations of each platform across different genomic contexts.

## Results

### Sequencing Quality Control

To evaluate the performance of our LRS dataset, we first conducted an analysis of the sequencing data quality and characteristics. We examined several key metrics, including sequencing yields, read length and quality distributions, alignment quality, sequencing depth, and flowcell and barcoding performance, to identify potential factors that may influence variant detection accuracy (**Figure 1**).

**Figure 1.**
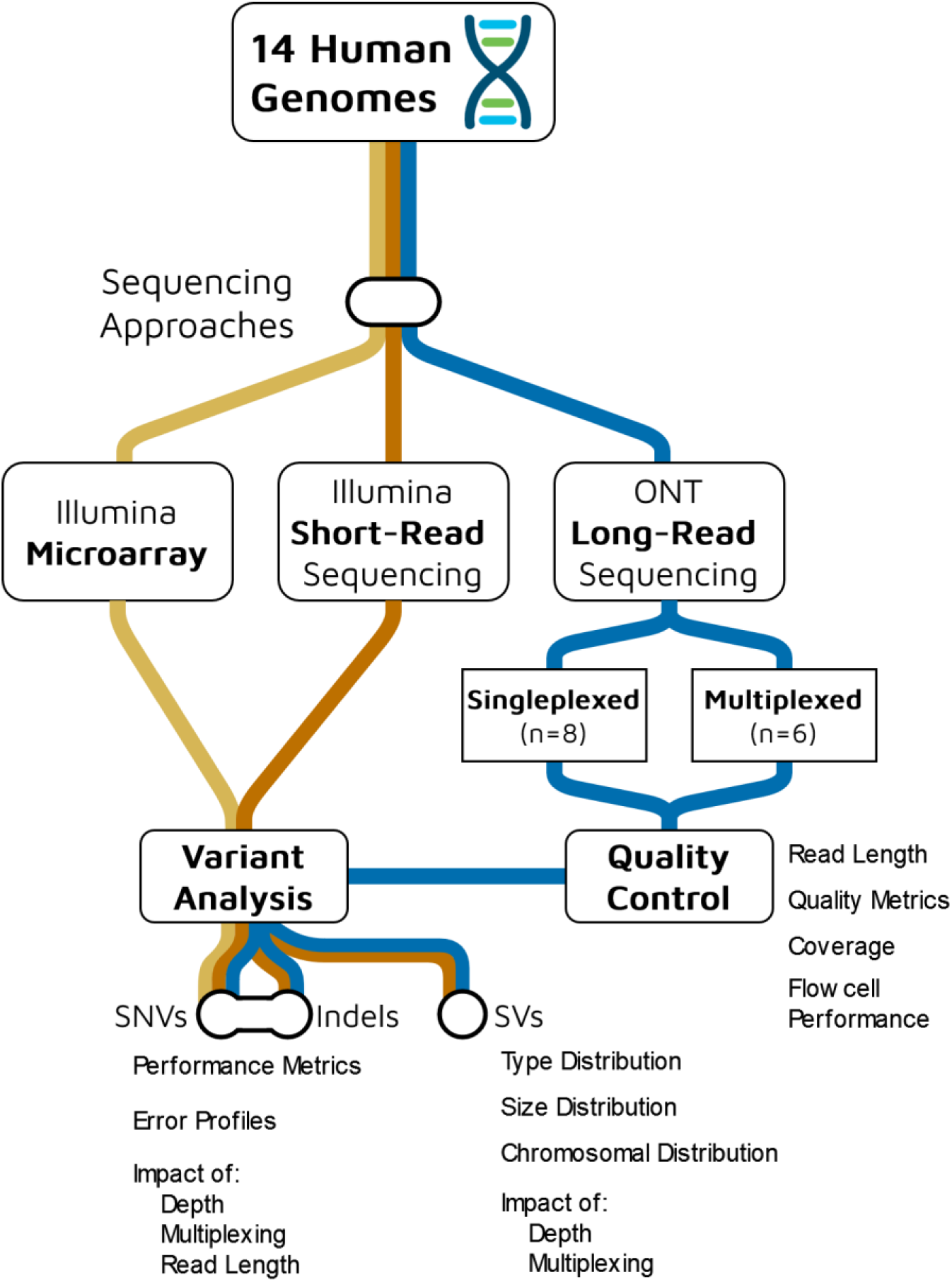
Experimental design and analysis workflow for comparing Oxford Nanopore Technologies (ONT) long-read sequencing (blue) with Illumina short-read sequencing (orange) and microarray (yellow) platforms across 14 human genomes.

### Sequencing Yields

Our analysis of multiplexed (two barcoded samples per flowcell) and singleplexed (one sample per flowcell) nanopore sequencing runs revealed significant differences in yield and read length characteristics between the two approaches.

In terms of sequencing yield, singleplexed samples consistently produced a higher number of reads and bases compared to multiplexed samples (**Figure 2B**). The mean number of reads for singleplexed samples was 9,932,389 (SD: 2,541,662), which is 118.48% higher than the mean of 4,546,234 (SD: 1,382,487) reads for multiplexed samples. Similarly, the mean number of bases for singleplexed samples (mean: 64,410,968,490; SD: 16,697,589,113) was 99.94% higher than that of multiplexed samples (mean: 32,214,905,852; SD: 5,350,881,127).5,350,881,127).

**Figure 2.**
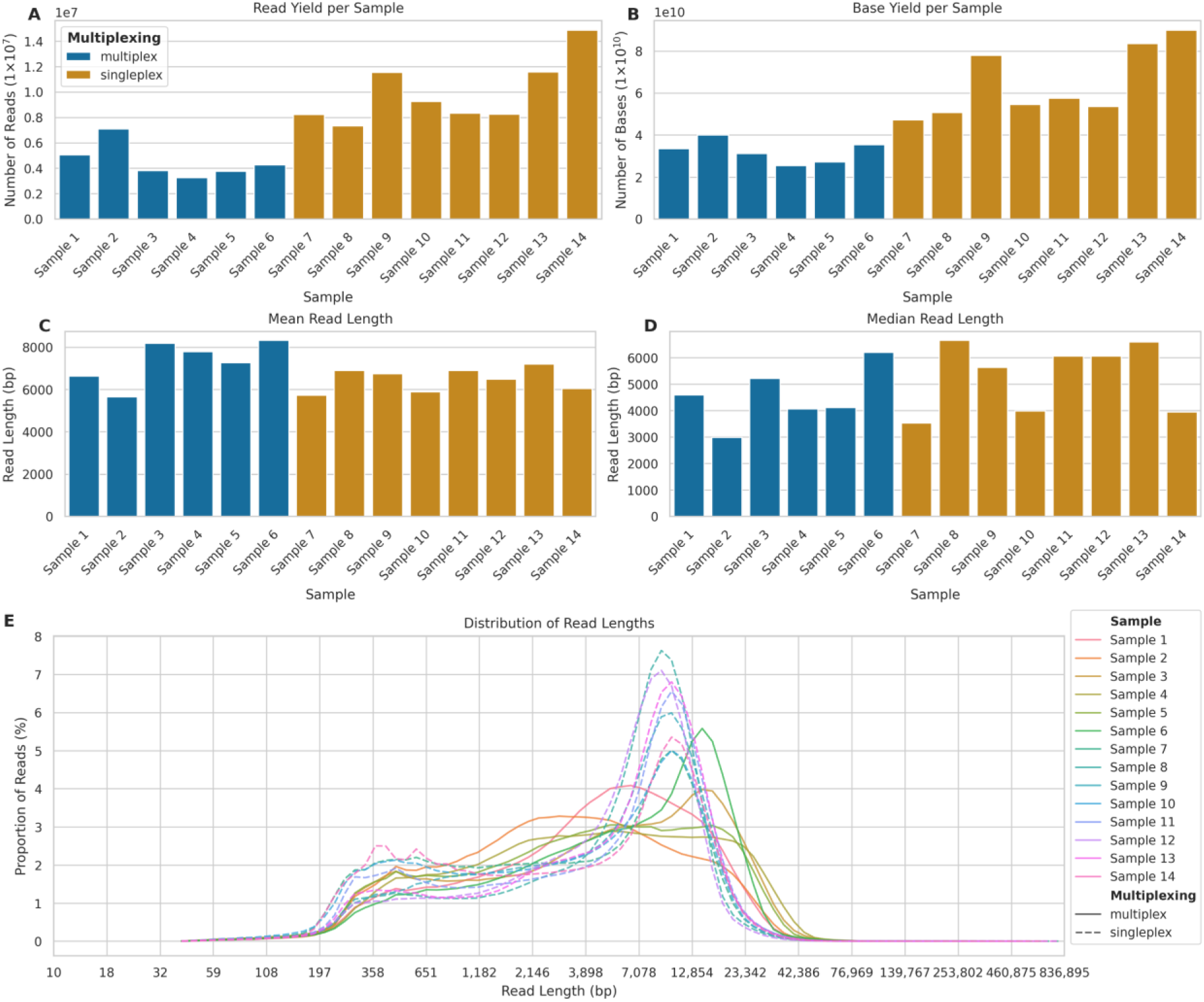
Comparison of read metrics between multiplexed and singleplexed ONT sequencing runs. **(A)** Number of reads per sample (×10^7^) for multiplexed (blue, Samples 1-6) and singleplexed (orange, Samples 7-14) runs. **(B)** Total bases per sample (×10^10^). (**C**) Mean and (**D**) Median read length in base pairs (bp). **(E)** Read length distribution across samples (solid lines: multiplexed; dashed lines: singleplexed) showing proportion of reads (%) in 100 logarithmic bins from 10 bp to 836,895 bp.

Interestingly, the read length distributions showed a different pattern (**Figure 2D**). While the mean read length for multiplexed samples (mean: 7,086 bp; SD: 7,668) was 8.48% higher than that of singleplexed samples (mean: 6,485 bp; SD: 6,155), the median read length for singleplexed samples (5,369 bp) was 27.08% higher than for multiplexed samples (4,225 bp). The distribution of read lengths across all samples (⍰⍰**Figure 2** Notably, all singleplexed samples exhibited a clear bimodal distribution of read lengths, characterised by two distinct peaks: one at shorter read lengths (approximately 400 bp) and another at longer read lengths (approximately 10,000 bp). In contrast, the multiplexed samples showed more variable distribution patterns. While some multiplexed samples displayed a bimodal distribution similar to the singleplexed samples, although with less pronounced peaks, other multiplexed samples exhibited only a single peak, typically at shorter read lengths (around 1,000-5,000 bp). In contrast, the multiplexed samples showed more variable distribution patterns. While some multiplexed samples displayed a bimodal distribution similar to the singleplexed samples, although with less pronounced peaks, other multiplexed samples exhibited only a single peak, typically at shorter read lengths (around 1,000-5,000 bp).

This difference in distribution patterns suggests that singleplexing consistently produced two distinct populations of reads: a shorter fraction and a longer fraction. Multiplexing, on the other hand, appeared to be less consistent in generating the longer read fraction, with some samples failing to produce the second peak at higher read lengths. This might explain why the larger number of reads per flowcell produced in singleplex did not translate into a larger number of genotyped bases.

### Read and Alignment Quality

Both read quality and alignment metrics showed minimal differences between multiplexed and singleplexed samples. Base quality distribution (**Figure 3A**) showed a similar pattern across all samples, with peaks at quality scores 17-22. The mean base quality for singleplexed samples (mean: 20.64; SD: 4.70) was marginally higher than that of multiplexed samples (mean: 20.44; SD: 4.69), representing a small 0.95% increase. Similarly, the median base quality for singleplexed samples (20.60) was slightly higher than multiplexed samples (20.34), a 1.24% increase.

**Figure 3.**
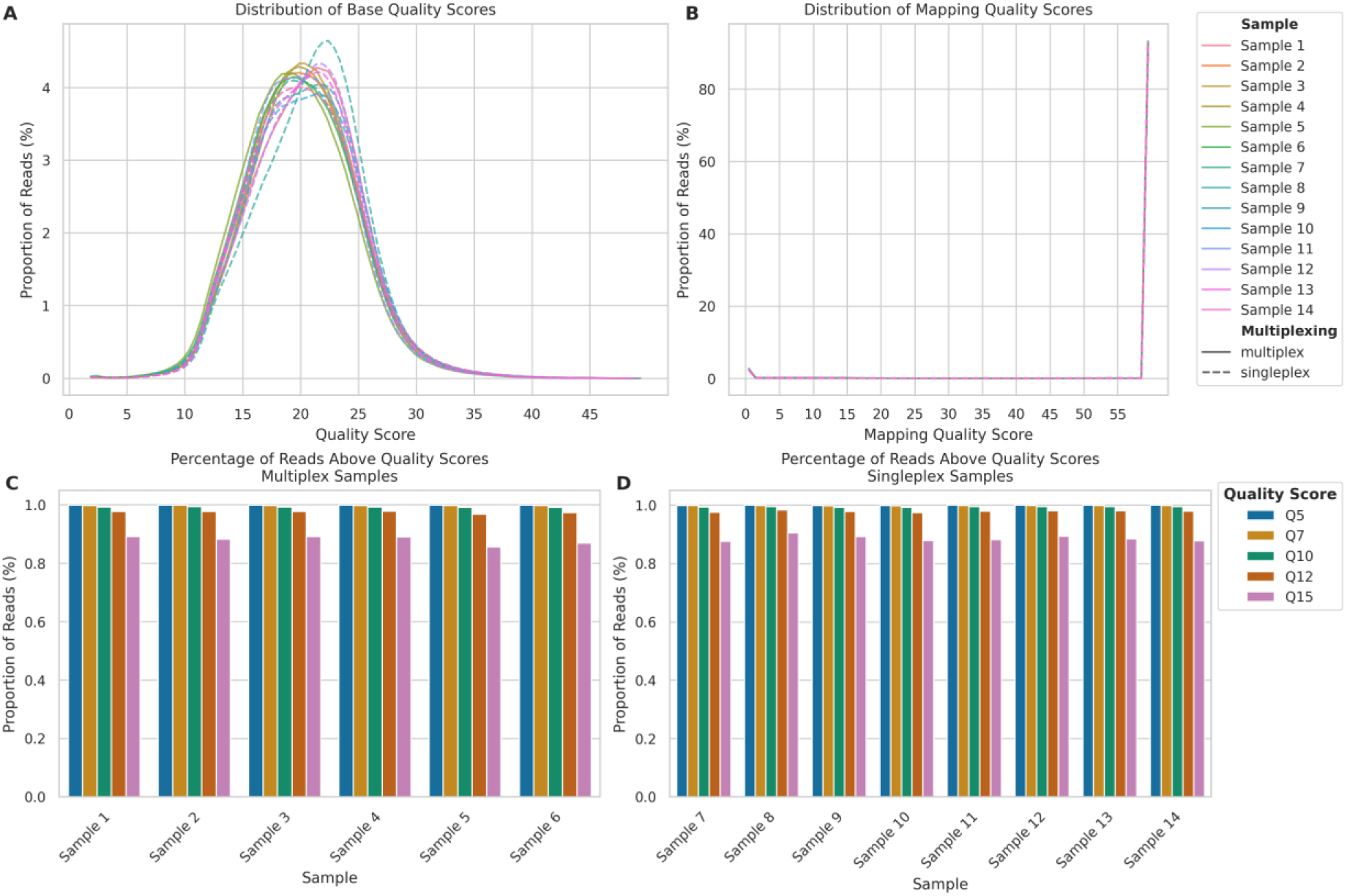
Comparison of sequencing quality metrics between multiplexed and singleplexed ONT sequencing runs. **(A)** Read quality score distributions for multiplexed (Samples 1-6, solid lines) and singleplexed samples (Samples 7-14, dashed lines) calculated by the Dorado basecaller. **(B)** Mapping quality score distributions, calculated by minimap2. **(C)** Percentage of reads above quality thresholds (Q5-Q15) for multiplexed Samples 1-6. **(D)** Percentage of reads above quality thresholds for singleplexed Samples 7-14.

The mapping quality distribution (**Figure 3B**) demonstrated that the vast majority of reads across all samples mapped with a quality score of 60. This pattern was nearly identical between singleplexed and multiplexed samples, with mean mapping qualities of 56.71 (SD: 12.67) and 56.68 (SD: 12.75), respectively, representing a negligible difference lower than 0.01%. The median mapping quality was 60 for both approaches, indicating no practical difference in mapping performance.

Both multiplexed and singleplexed samples showed a similar trend on the proportion of reads exceeding various quality score thresholds (Q5, Q7, Q10, Q12, and Q15) (**Figure 3C**, **D**). While the percentage of reads decreased as the quality threshold increased, at the highest threshold of Q15 all samples retained over 85% of their reads, suggesting multiplexing can be employed without significantly compromising data quality.

### Depth of Coverage

Focusing on sequencing depth of coverage per chromosome and across the whole genome, we observed differences between singleplexed and multiplexed samples in line with our expectations. Firstly, per-chromosome sequencing depth (**Figure 4A**) showed consistent patterns across all chromosomes, with singleplexed samples (mean: 19.01; SD: 5.98) exhibiting higher depth than multiplexed samples (mean: 9.45; SD: 2.33), even though singleplexed samples demonstrated greater variability. The mean whole genome depth (**Figure 4B**) mirrored the per-chromosome trends. Singleplexed samples achieved a mean depth of 19.84 (SD: 5.22), while multiplexed samples reached 9.84 (SD: 1.58), indicating a 101.63% increase in the mean depth of singleplexed samples. The median depths were 17.20 and 9.93 for singleplexed and multiplexed samples, respectively, showing a 73.30% increase. The consistent depth patterns across chromosomes show that ONT LRS maintains relatively uniform coverage across the genome in both settings, with expected drops in depth for sex chromosomes.

**Figure 4.**
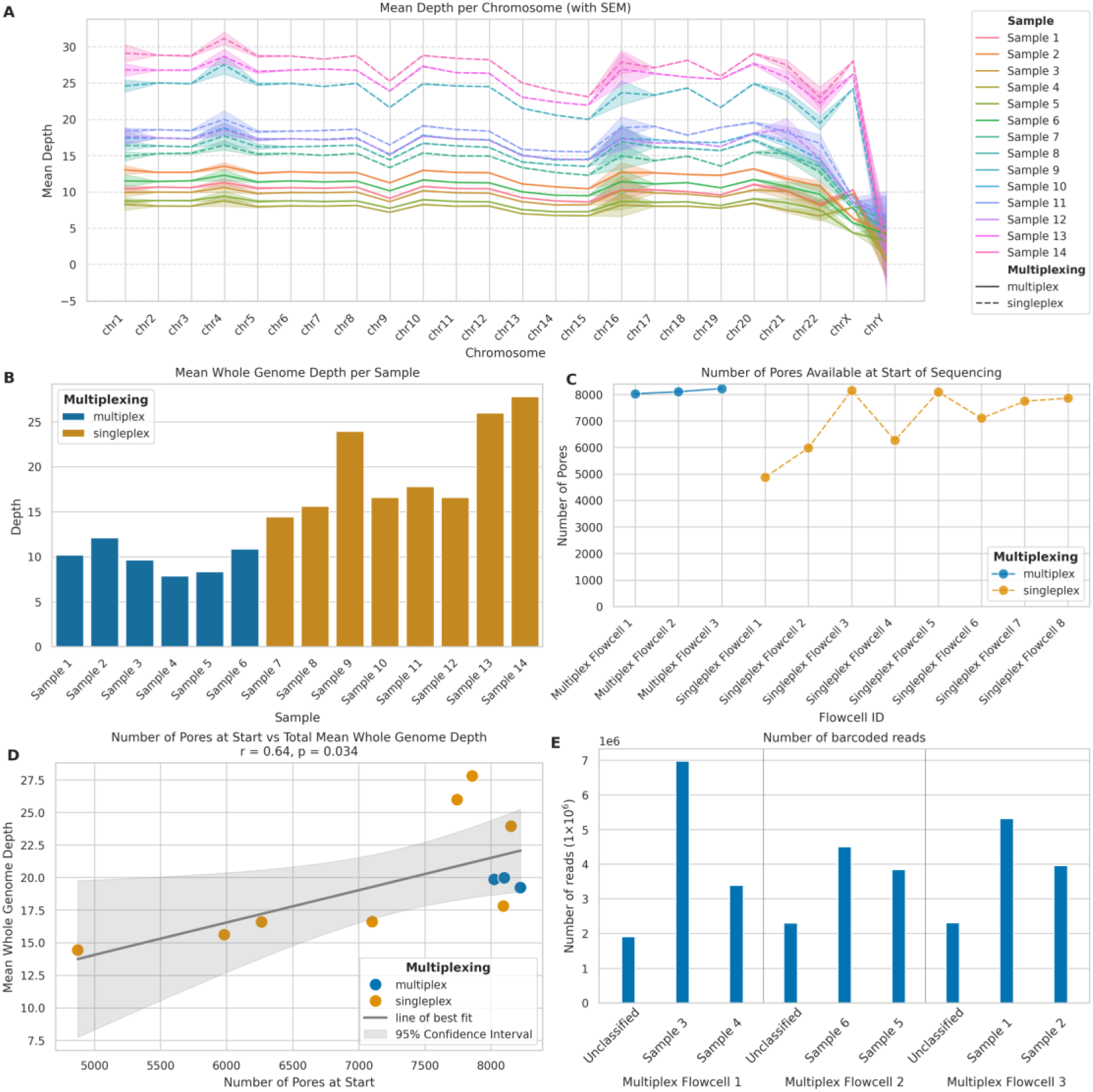
Comparison of multiplexed and singleplexed ONT sequencing performance. **(A)** Mean sequencing depth per chromosome for multiplexed (solid lines) and singleplexed (dashed lines) samples. Shaded areas: standard error of the mean (SEM). **(B)** Mean whole genome sequencing depth per sample for multiplexed (blue) and singleplexed (orange) runs. **(C)** Initial pore availability across flowcells for multiplexed (blue) and singleplexed (orange) runs. **(D)** Correlation between initial pore count and mean whole genome depth. Blue: multiplexed; orange: singleplexed. Solid line: linear regression; shaded area: 95% CI. **(E)** Read distribution across barcoded samples and unclassified reads in multiplexed flowcells.

### Flowcell and Barcoding Quality

We analysed the initial nanopore availability in the flowcells we used for sequencing to investigate its impact on sequencing performance and depth of coverage obtained. There were distinct differences in the number of pores available at the start of sequencing (**Figure 4C**), with multiplexed flowcells consistently having a higher number of starting pores (mean: 8,116.00; SD: 100.34) compared to singleplexed flowcells (mean: 7,008.75; SD: 1,190.38), which exhibited greater variability in starting pore numbers. This occurred because the best quality flowcells were selected for multiplexing to yield the best results possible for this approach. The relationship between initial pore count and total mean whole genome depth (**Figure 4D**) revealed a strong positive correlation (Pearson’s R-value: 0.64, p-value: 0.034). The linear regression analysis yielded a slope of 0.0025, which indicates that, for every additional 1,000 starting pores, we can expect an increase of approximately 2.5× in mean whole genome depth. This suggests that initial pore availability is a significant factor in determining sequencing depth yields.

Examining barcoded read distribution in multiplexed samples allowed us to assess the efficiency of the barcoding process and explain the differences in depth yields between multiplexed and singleplexed flowcells with similar number of nanopore availability (**Figure 4E**). Across the three multiplexed flowcells, we observed some variation in the number of reads assigned to the barcodes of interest and unclassified reads. To quantify this variation, we calculated the Coefficient of Variation (CV) for each flowcell: Multiplex Flowcell 1 showed the highest variation (CV: 63.78%), while Multiplex Flowcell 3 exhibited a CV of 38.91%; in contrast, Multiplex Flowcell 2 demonstrated the most consistent read distribution (CV: 31.77%). Importantly, an average of 19.04% (SD: 3.15%) of reads remained unclassified across all multiplexed flowcells. This level of variation within flowcells, and substantial proportion of unclassified reads indicates there is room for efficiency improvement in the barcoding process and, if not optimised, multiplexing might result in a lower total number of genotyped bases per flowcell.

### Single Nucleotide Variants Benchmark

We compared the performance of the ONT and Illumina sequencing platforms for SNV detection with the VCFeval tool of RTG tools (37, 38). Given their high accuracy in genotyping SNVs, we used the Illumina microarray datasets as the gold standard, stratified by high complexity regions (i.e., excluding the Tandem Repeats and Homopolymers regions of GIAB v3.4 (33)) and low complexity genomic regions (only including the aforementioned sites). In addition, only the sites present in the SNP arrays were retained in SRS and LRS data for comparison. For ONT LRS, we analysed 9,553,609 variants in high complexity regions and 491,665 variants in low complexity regions. For SRS, we analysed 9,466,960 variants in high complexity regions and 482,332 variants in low complexity regions. **Figure 5**, **B** shows the precision, sensitivity, and F-measure for both technologies in high-and low-complexity regions of the genome..

**Figure 5.**
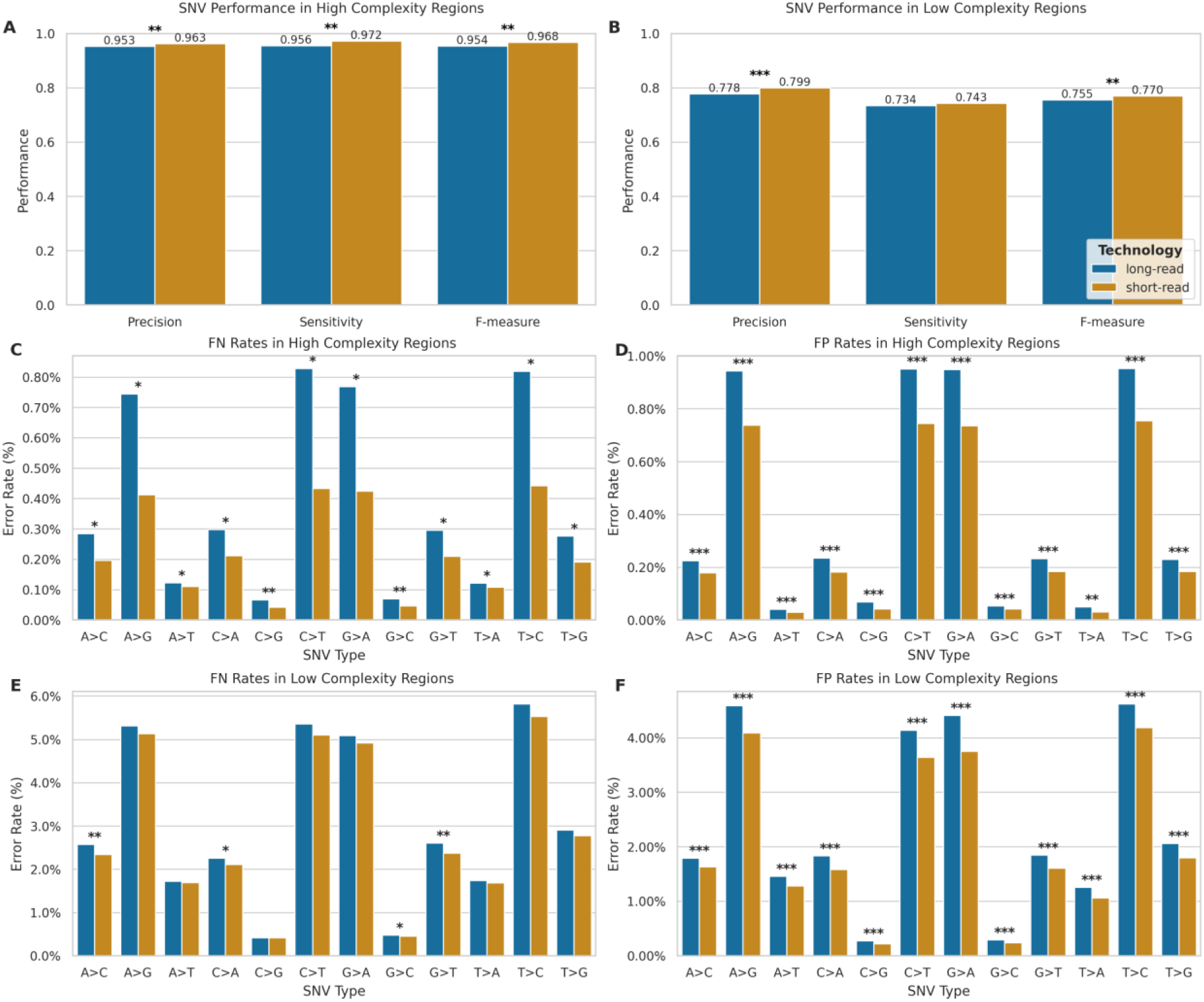
Comparison of SNV detection performance between ONT LRS and SRS. **(A, B)** Performance metrics (precision, sensitivity, F-measure) in high (A; LRS: n=9,553,609; SRS: n=9,467,517) and low complexity regions (B; ONT: n=491,665; Illumina: n=482,332). Blue: long-read; orange: short-read. **p < 0.01, ***p < 0.001. **(C-F)** SNV-specific error rates showing false negatives (C, E) and false positives (D, F) in high (C, D) and low complexity regions (E, F). X-axis indicates nucleotide substitutions. *p < 0.05, **p < 0.01, ***p < 0.001.

In high complexity regions, SRS maintains an advantage over LRS technology, with statistically significant differences (p < 0.01) in mean precision (SRS: 0.963, SD = 0.004; LRS: 0.953, SD = 0.010), sensitivity (SRS: 0.972, SD = 0.001; LRS: 0.956, SD = 0.021), and F-measure (SRS: 0.968, SD = 0.002; LRS: 0.954, SD = 0.015) compared to LRS. However, the performance slightly shifted in low complexity regions. While SRS still exhibited higher mean precision (SRS: 0.799, SD = 0.005; LRS: 0.778, SD = 0.013, p < 0.001) and F-measure (SRS: 0.770, SD = 0.003; LRS: 0.755, SD = 0.016, p < 0.01), the gap in performance narrowed considerably for mean sensitivity (SRS: 0.743, SD = 0.002; LRS: 0.734, SD = 0.019), with no statistically significant difference (p > 0.05) between platforms. On the whole, the performance of both technologies degrades in low complexity regions, but to different degrees, with a more pronounced performance drop for SRS.

To further elucidate the performance of each technology, we conducted an in-depth analysis of SNV error rates, stratified by complexity and error type (**Figure 5-F**), revealing distinct error profiles for SRS and LRS across different SNV types. In high complexity regions, LRS exhibited significantly higher error rates across all SNV types for both false positives (FP) and false negatives (FN) (p < 0.05 for all comparisons, ⍰**Figure 5D**). The most pronounced differences in performance were observed in transitions (A>G, G>A, C>T, and T>C). The error profile slightly shifted in low complexity regions (⍰⍰**Figure 5**). While for FPs, LRS maintained significantly higher error rates across all SNV types (p < 0.001), with the largest differences seen again in transitions, for FNs LRS demonstrated significantly higher error rates for transversions A>C, C>A, and G>C, as well as the transition C>T (p < 0.001), while most other SNV types showed no significant differences between technologies. with the largest differences seen again in transitions, for FNs LRS demonstrated significantly higher error rates for transversions A>C, C>A, and G>C, as well as the transition C>T (p < 0.001), while most other SNV types showed no significant differences between technologies.

While SRS generally maintains an advantage in high complexity regions and in minimising false positives, LRS shows competitive performance in detecting certain SNV types within low complexity regions, particularly in terms of false negative rates. The observed differences in error profiles between the two technologies likely stem from their fundamental sequencing approaches. SRS’s short-read methodology may struggle with repetitive regions due to mapping ambiguities, while LRS’s long-read approach can span these regions but introduces more base-calling errors, particularly for transitions.

### Indels Benchmark

To evaluate the performance of LRS for indel detection using VCFeval, we utilised the SRS dataset as the gold-standard, as SRS has demonstrated superior performance in indel detection with F1 scores of up to 0.99 for state-of-the-art variant calling pipelines (39).

Our results showed a marked disparity in performance between high complexity and low complexity genomic regions (**Figure 6A**, **B**). On one hand, in high complexity areas, LRS exhibited robust performance with a mean precision of 0.786 (SD = 0.067), sensitivity of 0.905 (SD = 0.035), and F-measure of 0.841 (SD = 0.053). Conversely, in low complexity regions, we observed a significant decline in performance metrics, with mean precision, sensitivity, and F-measure dropping to 0.395 (SD = 0.094), 0.462 (SD = 0.039), and 0.423 (SD = 0.069), respectively.

**Figure 6.**
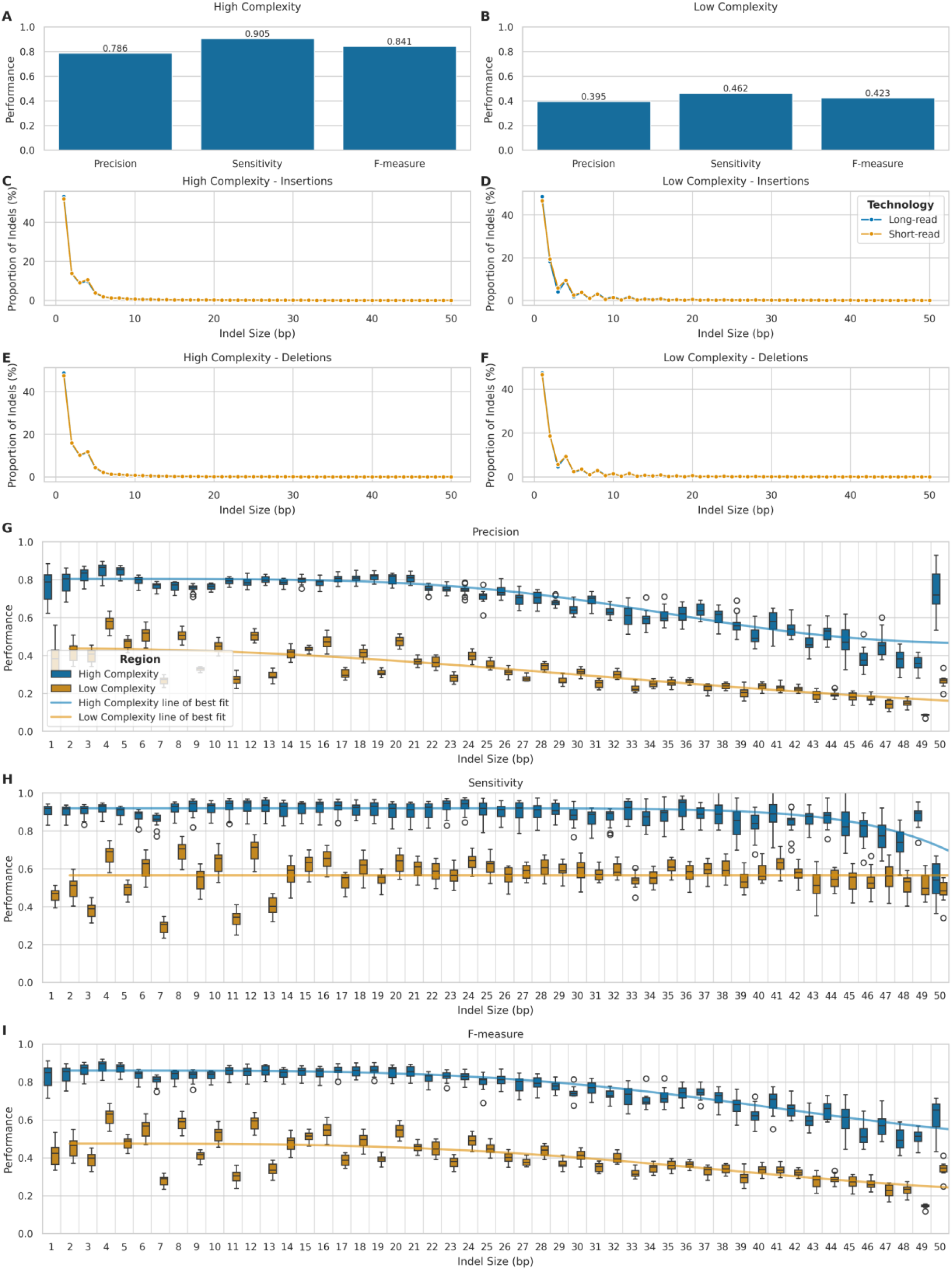
Comparison of Indel detection performance between ONT LRS and SRS. (**A-B**) Precision, sensitivity, and F-measure for ONT indel detection in high (A) and low (B) complexity regions. (**C-F**) Size distribution of indels detected by ONT and Illumina: insertions (C,D) and deletions (E,F) in high (C,E) and low (D,F) complexity regions. (**G-I**) ONT indel calling metrics by indel length (1-50 bp) in high (blue) and low (orange) complexity regions: precision (G), sensitivity (H), and F-measure (I).

We further analysed the size distribution of insertions and deletions detected by both LRS and SRS platforms in high and low complexity genomic regions (**Figure 6C-F**), to ascertain potential biases in the detection of indels of various sizes. We classified the variants into insertions and deletions, within their respective complexity regions. Indeed, both technologies performed comparably in detecting small indels within high complexity genomic regions (**Figure 6C**, **E**), with similar size distributions peaking at 1-2 bp and rapidly decreasing for larger sizes.

To quantify these observations, we calculated summary statistics for indel sizes detected by each platform. The Kolmogorov-Smirnov test revealed minimal differences between platforms in these regions (KS statistic: 0.013 for insertions, 0.013 for deletions). However, in low complexity regions (**Figure 6D**, **F**) LRS demonstrated a higher propensity for detecting larger indels, particularly around 8-12 bp. Though this disparity was slightly more pronounced, the effect sizes were still small (KS statistic: 0.028 for insertions, 0.012 for deletions). All differences were statistically significant (p-values < 2.1×10^-116^ after FDR correction).

To further investigate the performance characteristics of LRS across different indel lengths, we stratified the analysis by indel size categories and complexity regions (**Figure 6G - I**). In high complexity regions, LRS demonstrated robust performance for smaller indels (1-5bp), which represented the majority of variants (89.5%), achieving high precision (median=0.826, IQR=0.071), sensitivity (median=0.918, IQR=0.040), and F-measure (median=0.869, IQR=0.066). Performance remained stable for medium-sized indels (6-20bp), with only modest decreases in precision (median=0.772-0.798) and sensitivity (median=0.885-0.932). However, there was a notable decline in performance for larger indels (21-50bp), particularly in precision (median=0.624, IQR=0.184) while maintaining relatively high sensitivity (median=0.885, IQR=0.097).

In low complexity regions, we observed substantially lower performance across all size categories. For the predominant 1-5bp small indels (83.5% of variants), precision (median=0.445, IQR=0.116) and sensitivity (median=0.483, IQR=0.109) were markedly reduced compared to high complexity regions. Performance further deteriorated with increasing indel size, with the largest indels (21-50bp) showing the lowest precision (median=0.250, IQR=0.093), maintaining moderate sensitivity (median=0.571, IQR=0.085). Stretched exponential regression analysis revealed strong relationships between indel size and performance metrics in high complexity regions (R² = 0.849, 0.670, and 0.913 for precision, sensitivity, and F-measure respectively), while relationships were weaker or absent in low complexity regions (R² = 0.690, 0.000, and 0.575 respectively). Therefore, this confirms that indel size significantly impacts variant calling accuracy, with the effect being more pronounced in high complexity regions.

This size-dependent performance pattern indicates that while LRS technology maintains reliable detection of small to medium-sized indels in high complexity regions, its accuracy is substantially compromised in low complexity regions regardless of indel size. The particularly poor performance for larger indels in low complexity regions highlights the ongoing challenges in accurately sequencing and analysing repetitive or structurally complex genomic areas. However, when considering the inherent limitations of our benchmarking approach, it is important to interpret these results with caution. While we used Illumina SRS as the gold standard for this comparison, it is important to note that SRS itself has limitations, particularly in sequencing repetitive regions. The increasing error rates with increasing indel size could partially reflect the limitations of SRS in detecting larger indels, especially in repetitive regions. Indeed, some of the false positives in LRS, particularly for larger indels in low complexity regions, may actually represent true variants that were missed by SRS.

### Impact of multiplexing on variant calling

We compared the performance of variant calling, for both SNVs and indels in high complexity and low complexity regions, for multiplexed and singleplexed samples, to determine the effect of multiplexing on the accuracy of variant calling.

For SNVs in high complexity regions (**Figure 7A**), singleplexed samples demonstrated marginally superior performance across all metrics. Indeed, mean precision (singleplex: 0.960, SD = 0.003; multiplex: 0.944, SD = 0.008), sensitivity (singleplex: 0.970, SD = 0.004; multiplex: 0.936, SD = 0.018), and F-measure (singleplex: 0.965, SD = 0.003; multiplex: 0.940, SD = 0.013) were all higher for singleplexed samples. A similar trend was evident in low complexity regions (**Figure 7B**), with higher mean precision (singleplex: 0.788, SD = 0.005; multiplex: 0.765, SD = 0.009), sensitivity (singleplex: 0.747, SD = 0.005; multiplex: 0.717, SD = 0.015), and F-measure (singleplex: 0.767, SD = 0.005; multiplex: 0.740, SD = 0.012) for singleplexed samples.

**Figure 7.**
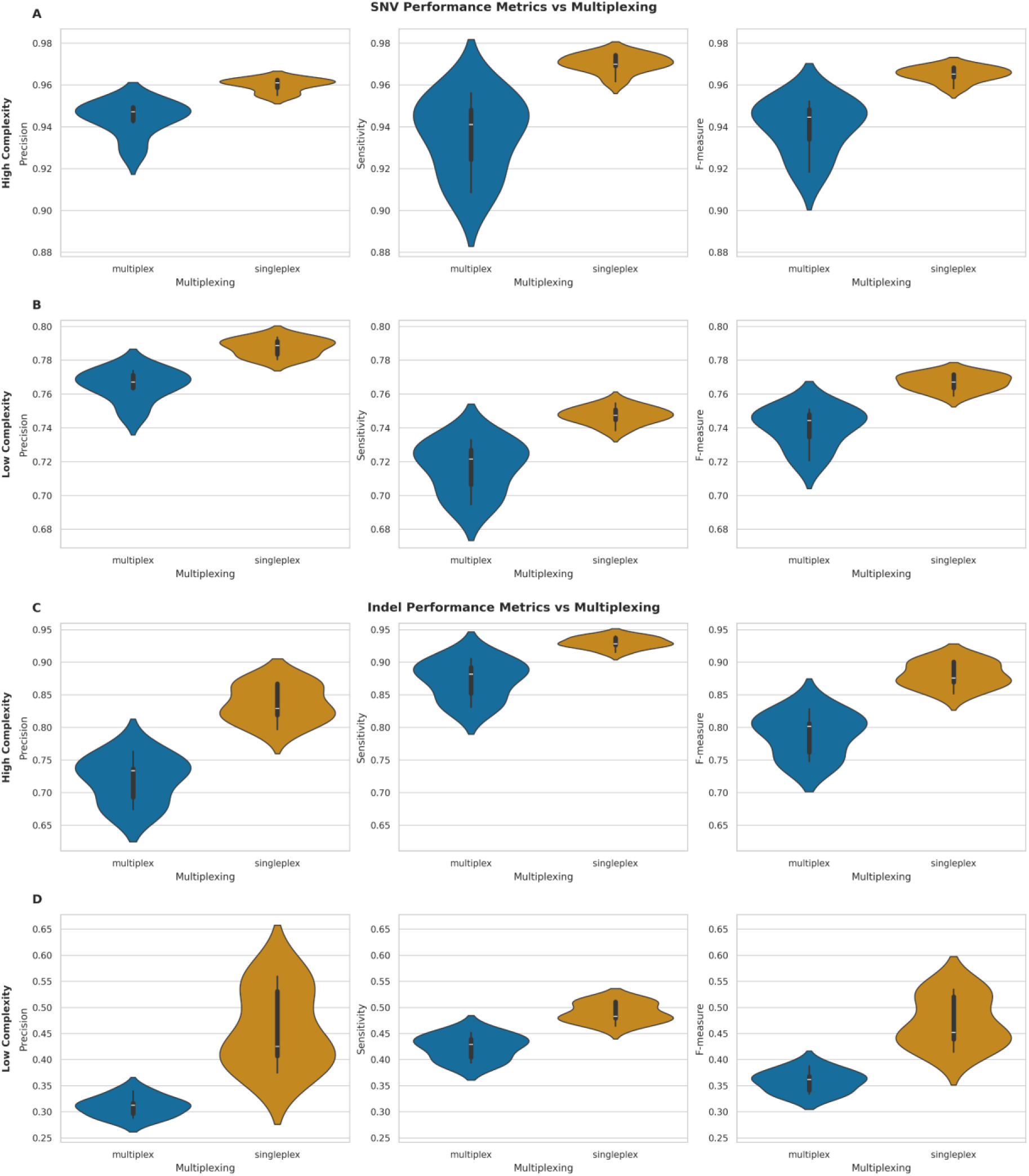
Performance metrics comparison between multiplexed and singleplexed samples for SNV and Indel detection. **(A-B)** Violin plots of Precision, Sensitivity, and F-measure for SNVs in high (A) and low (B) complexity genomic regions. **(C-D)** Violin plots of same metrics for Indels in high (C) and low (D) complexity regions. All plots compare multiplexed (blue) vs. singleplexed (orange) samples, with white bars indicating median values.

Indel calling exhibited more pronounced differences between multiplexing conditions. In high complexity regions (**Figure 7C**), singleplexed samples showed substantial improvements in mean precision (singleplex: 0.836, SD = 0.028; multiplex: 0.720, SD = 0.035), sensitivity (singleplex: 0.930, SD = 0.009; multiplex: 0.873, SD = 0.030), and F-measure (singleplex: 0.880, SD = 0.019; multiplex: 0.789, SD = 0.033). The disparity was even more evident in low complexity regions (**Figure 7D**), with higher mean precision (singleplex: 0.458, SD = 0.074; multiplex: 0.310, SD = 0.019), sensitivity (singleplex: 0.490, SD = 0.019; multiplex: 0.424, SD = 0.024), and F-measure (singleplex: 0.472, SD = 0.048; multiplex: 0.358, SD = 0.021) for singleplexed samples.

To separate the effects of multiplexing and sequencing depth, we performed an analysis of covariance (ANCOVA) (**Figure 8**). This analysis revealed that sequencing depth had a significant positive effect on all performance metrics (p < 0.001), with a particularly strong impact on indel calling (R² > 0.95). The independent effect of multiplexing, after controlling for depth, was considerably smaller and reached statistical significance only for indel precision in both high complexity (effect size = -0.021, p = 0.013) and low complexity regions (effect size = -0.008, p = 0.010). For SNV calling, the multiplexing effect was not statistically significant across any metric (p > 0.15), suggesting that the apparent performance differences were primarily driven by variations in sequencing depth rather than multiplexing itself.

**Figure 8.**
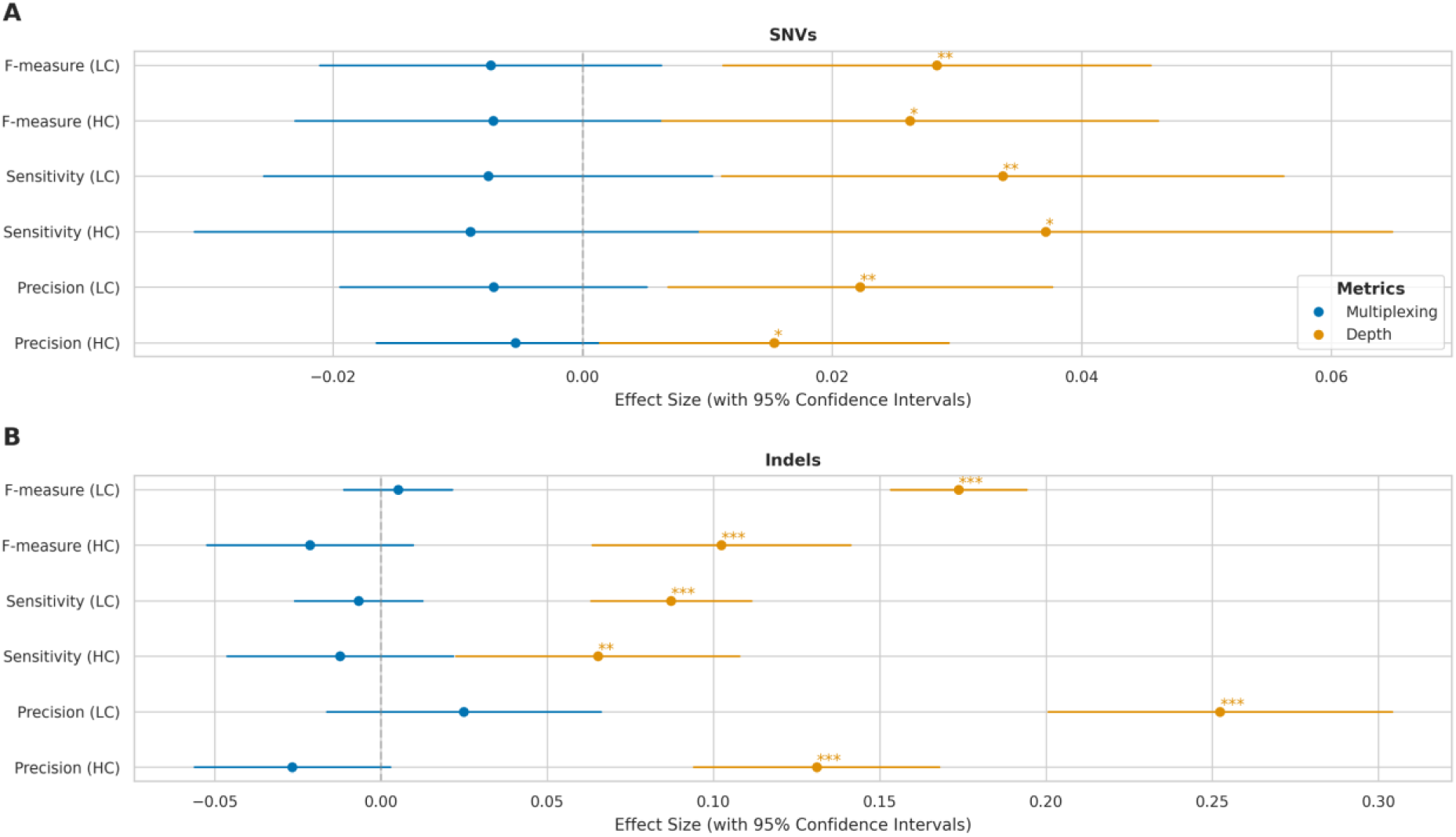
Multiplexing and sequencing depth effects on variant calling across genomic complexity regions. **(A)** Forest plot showing effect sizes with 95% confidence intervals for SNV detection metrics (Precision, Sensitivity, F-measure) in high complexity (HC) and low complexity (LC) regions. Blue: multiplexing effects (relative to singleplex); orange: log-transformed sequencing depth effects. **(B)** Corresponding effect sizes for Indel detection metrics. X-axis shows magnitude and direction of effects (negative values indicate decreased performance). Statistical significance: * p < 0.05, ** p < 0.01, *** p < 0.001.

Thus, while initial comparisons suggested that multiplexing has a modest impact on SNV calling performance and more significantly affects indel calling accuracy, particularly in low complexity regions, our subsequent analysis indicates that these effects are largely attributable to differences in sequencing depth rather than multiplexing per se. These findings suggest that maintaining adequate sequencing depth is more critical for variant calling performance than the choice between multiplexed and singleplexed approaches.

### Impact of sequencing depth on variant calling

To assess the influence of sequencing depth on variant calling performance, we analysed the relationship between whole genome mean depth and the variant calling performance metrics (precision, sensitivity, and F-measure) for both SNVs and indels in high complexity and low complexity regions.

For SNVs in high complexity regions (**Figure 9A**), we observed strong positive correlations between sequencing depth and all performance metrics (precision: r = 0.81, p = 4.94 ×10^- 4^; sensitivity: r = 0.81, p = 3.89×10^-4^; F-measure: r = 0.82, p = 3.42×10^-4^). Similar trends were observed for SNVs in low complexity regions (**Figure 9B**), with slightly stronger correlations (precision: r = 0.86, p = 6.92×10^-5^; sensitivity: r = 0.83, p = 2.06×10^-4^; F-measure: r = 0.86, p = 8.42×10^-5^).

**Figure 9.**
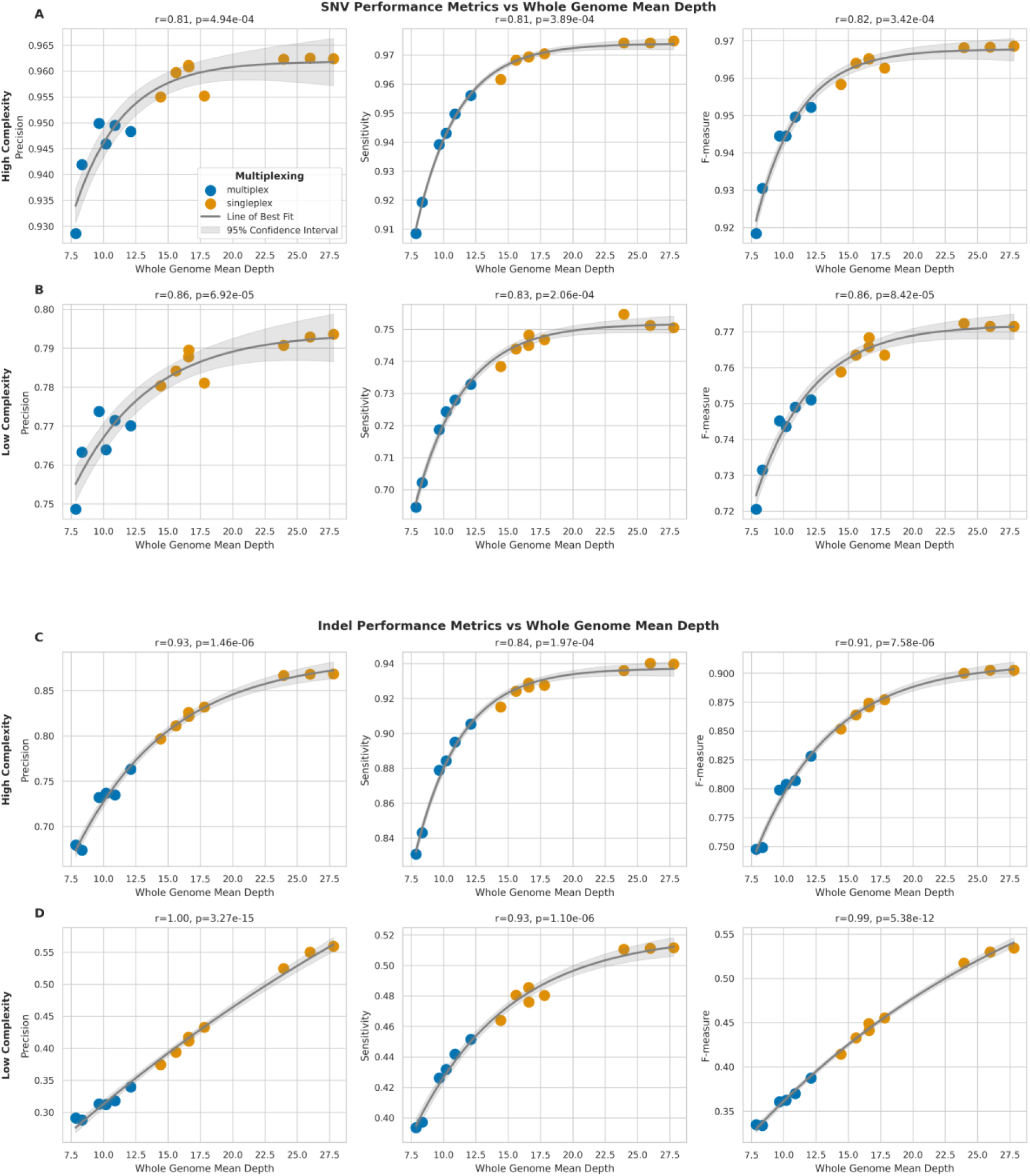
Impact of sequencing depth on SNV and indel detection performance across genomic regions. **(A-B)** Relationship between whole genome mean depth and performance metrics (Precision, Sensitivity, F-measure) for SNV detection in high (A) and low (B) complexity regions. **(C-D)** Relationship between whole genome mean depth and performance metrics for indel detection in high (C) and low (D) complexity regions. Grey curves represent asymptotic function fits with 95% confidence intervals (light grey). Pearson correlation coefficients (r) with p-values are shown.

Indel calling performance showed even stronger correlations with sequencing depth. In high complexity regions (**Figure 9C**), the correlations were highly significant for all metrics (precision: r = 0.93, p = 1.46×10^-6^; sensitivity: r = 0.84, p = 1.97×10^-4^; F-measure: r = 0.91, p = 7.58×10^-6^), but the strongest correlations were observed for indels in low complexity regions (**Figure 9D**) (precision: r = 1.00, p = 3.27×10^-15^; sensitivity: r = 0.93, p = 1.10×10^-6^; F-measure: r = 0.99, p = 5.38×10^-12^).

On the whole, the relationship between sequencing depth and performance metrics followed an asymptotic trend for all variant types and genomic regions, therefore suggesting that while increasing sequencing depth generally improves variant calling performance, the rate of improvement plateaus at higher depths. Additionally, the impact of sequencing depth is more pronounced for indel calling compared to SNV calling, particularly in low complexity regions.

### Impact of read length on variant calling

To investigate the relationship between read length and variant calling performance, we analysed the correlation between mean read length and the three performance metrics: precision, sensitivity, and F-measure, for both SNVs and indels, in high complexity and low complexity regions.

For SNVs in high complexity regions (**Figure 10A**), we observed moderate negative correlations between mean read length and all three performance metrics (precision: r = -0.43, p = 0.126; sensitivity: r = -0.51, p = 5.97×10^-2^; F-measure: r = -0.49, p = 7.35×10^-2^). Similar negative correlations were observed for SNVs in low complexity regions (**Figure 10B**) (precision: r = -0.41, p = 0.149; sensitivity: r = -0.50, p = 6.70×10^-2^; F-measure: r = - 0.47, p = 8.76×10^-2^).

**Figure 10.**
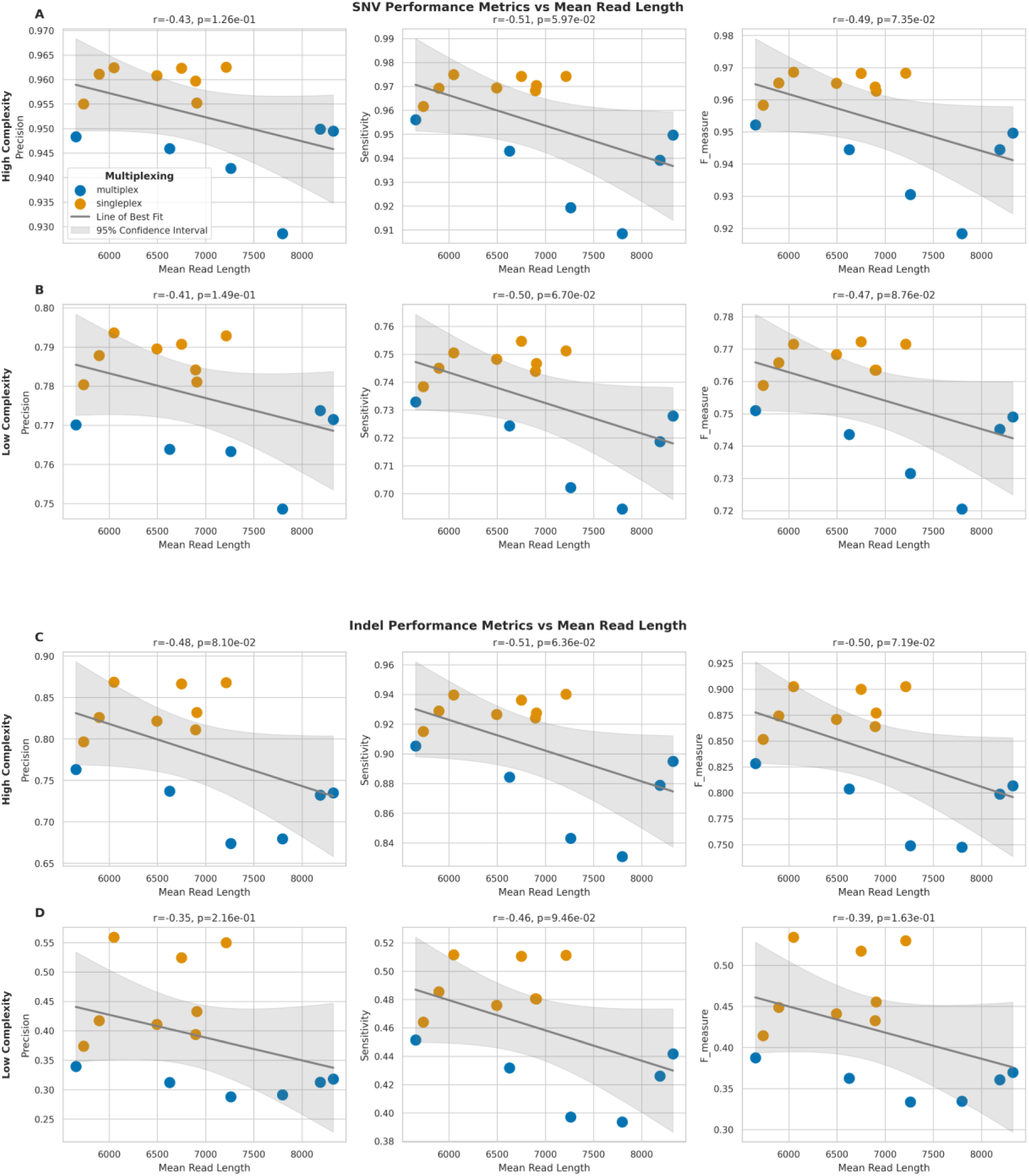
Impact of mean read length on SNV and indel detection performance across genomic regions. **(A-B)** Relationship between mean read length and performance metrics (Precision, Sensitivity, F-measure) for SNV detection in high (A) and low (B) complexity regions. **(C-D)** Relationship between mean read length and performance metrics for indel detection in high (C) and low (D) complexity regions. Blue/orange dots represent multiplex/singleplex samples, respectively. Blue lines show best fit with 95% confidence intervals (shaded). Pearson correlation coefficients (r) with p-values are shown.

Moreover, indel calling performance also showed comparable negative correlations with mean read length. In high complexity regions (**Figure 10C**), the correlations were slightly stronger than those observed for SNVs (precision: r = -0.48, p = 8.10×10^-2^; sensitivity: r = -0.51, p = 6.36×10^-2^; F-measure: r = -0.50, p = 7.19×10^-2^), and, for indels in low complexity regions (**Figure 10D**), the correlations were also negative and statistically significant (precision: r = -0.35, p = 0.216; sensitivity: r = -0.46, p = 9.46×10^-2^; F-measure: r = -0.39, p = 0.163).

Therefore, these results suggest a trend toward longer read lengths being associated with a moderate decrease in variant calling performance for both SNVs and indels, across both high and low complexity regions, although most correlations did not reach statistical significance.

### Structural Variant Calling Evaluation

To evaluate the performance of ONT LRS and Illumina SRS platforms in detecting structural variants (SVs), we conducted a consensus analysis with SURVIVOR (40) to merge consensus SV calls from both technologies. SV calls were considered in consensus if they met the following criteria: (1) located within 500 base pairs of each other, (2) detected in both VCF files for a given sample, (3) classified as the same SV type, (4) identified on the same strand, and (5) exhibiting a size difference of no more than 30% between platforms.

Overall, ONT LRS identified 2.86 times more SVs compared to Illumina SRS across all samples (**Figure 11A**). However, the consensus between the two platforms was relatively low, with an average of 20.61% (SD = 1.56%) of LRS calls and 57.87% (SD = 9.73%) of SRS calls being concordant. Furthermore, regarding SV size distribution (**Figure 11B**), LRS demonstrated a broader range of SV sizes, particularly in detecting larger variants, while Illumina SRS showed a bias towards detecting smaller SVs. In fact, LRS detected SVs ranging from 12 bp to 129,371,498 bp, with a mean size of 3,527.59 bp (SD = 482,253.95 bp) and a median of 80 bp. In contrast, SRS identified SVs between 2 bp and 6,064 bp, with a mean size of 165.62 bp (SD = 164.47 bp) and a median of 96 bp. The markedly larger maximum SV size (129 Mb for LRS vs 6 Kb for SRS) and higher SD observed in LRS data highlight its superior capability in detecting large-scale genomic rearrangements, which are often challenging to identify with SRS.

**Figure 11.**
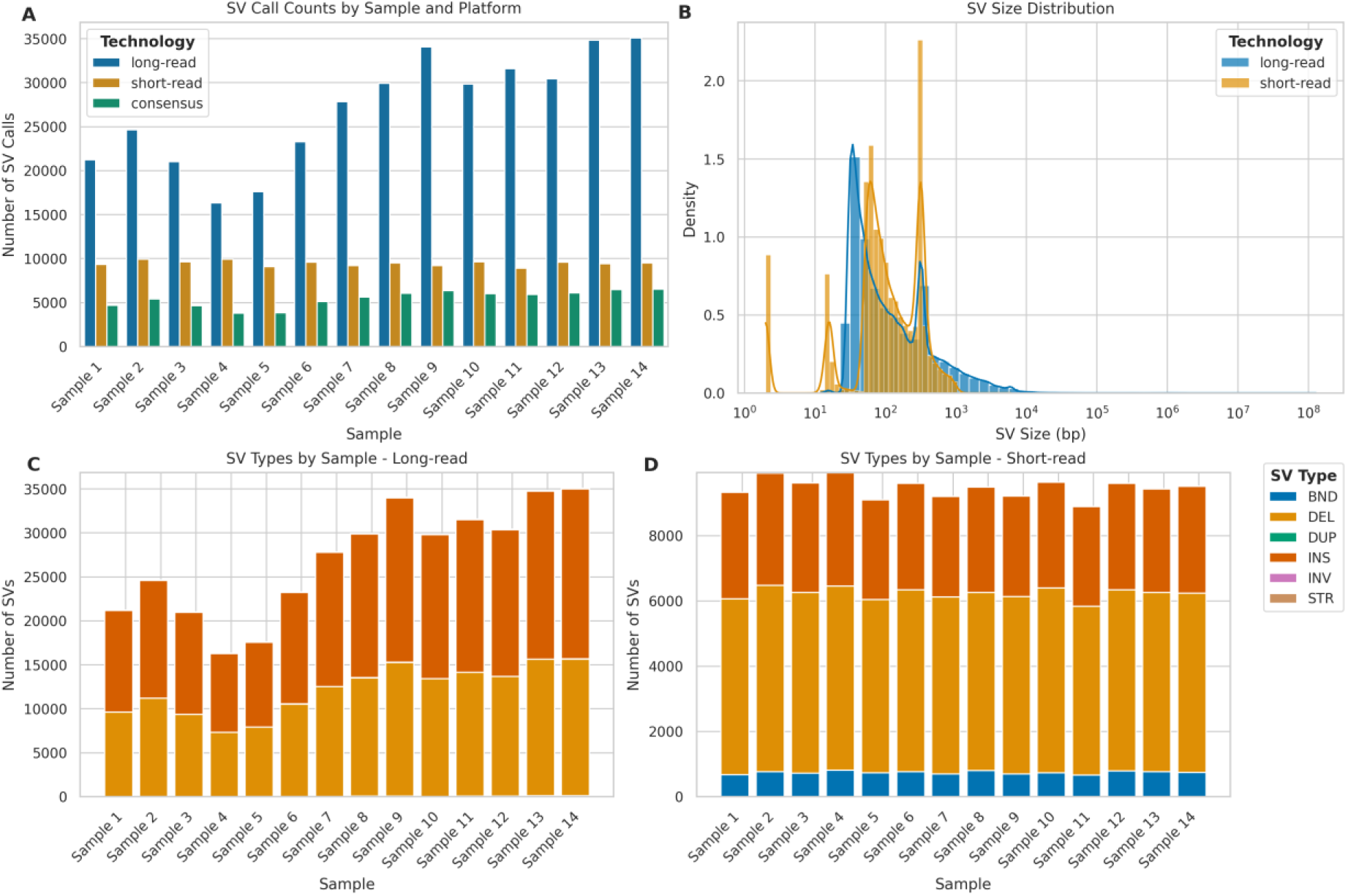
Comparative analysis of structural variant (SV) detection using long-read (ONT) and short-read (Illumina) sequencing platforms. **(A)** SV calls per sample: long-read (blue), short-read (orange), and consensus calls (green). **(B)** SV size distribution (log scale) for long-read (blue) and short-read (orange) platforms. **(C)** Distribution of SV types; BND (Breakend), DEL (Deletion), DUP (Duplication), INS (Insertion), INV (Inversion), STR (Short Tandem Repeat) detected by long-read sequencing across samples. **(D)** Distribution of SV types detected by short-read sequencing across samples.

To further investigate the nature of these SV calls, we analysed the distribution of SV types for each sequencing platform (**Figure 11C**, **D**). For LRS, insertions were the most frequently detected (mean = 14,784.64, SD = 3,352.94, per sample), followed closely by deletions (mean = 12,074.57, SD = 2,675.39). SRS, however, detected deletions most frequently (mean = 5,491.57, SD = 139.84), followed by insertions (mean = 3,230.07, SD = 127.40). Notably, SRS detected a higher number of breakends (BND) (mean = 741.21, SD = 44.07) compared to LRS (mean = 53.14, SD = 35.98), while LRS showed the capability to detect STRs (mean = 25.64, SD = 8.21).

The size distribution of different SV types further showed distinct patterns between LRS and SRS platforms. For insertions (**Figure 12A**), both technologies showed multimodal distributions, with LRS capturing a broader range of sizes. SRS exhibited a trimodal distribution with a sharp peak at smaller insertion sizes (< 10 bp), a second peak around 100 bp, and a third, broader peak centred approximately at 300 bp. In contrast, LRS demonstrated a bimodal distribution with peaks at approximately 300 bp and 6 kb. Deletion size distributions (**Figure 12B**) showed similar bimodal patterns for both platforms, with LRS again exhibiting a wider range. SRS deletions were predominantly clustered below 1 kb, while LRS effectively captured deletions spanning several orders of magnitude, from 50 bp to 10 Mb.

**Figure 12.**
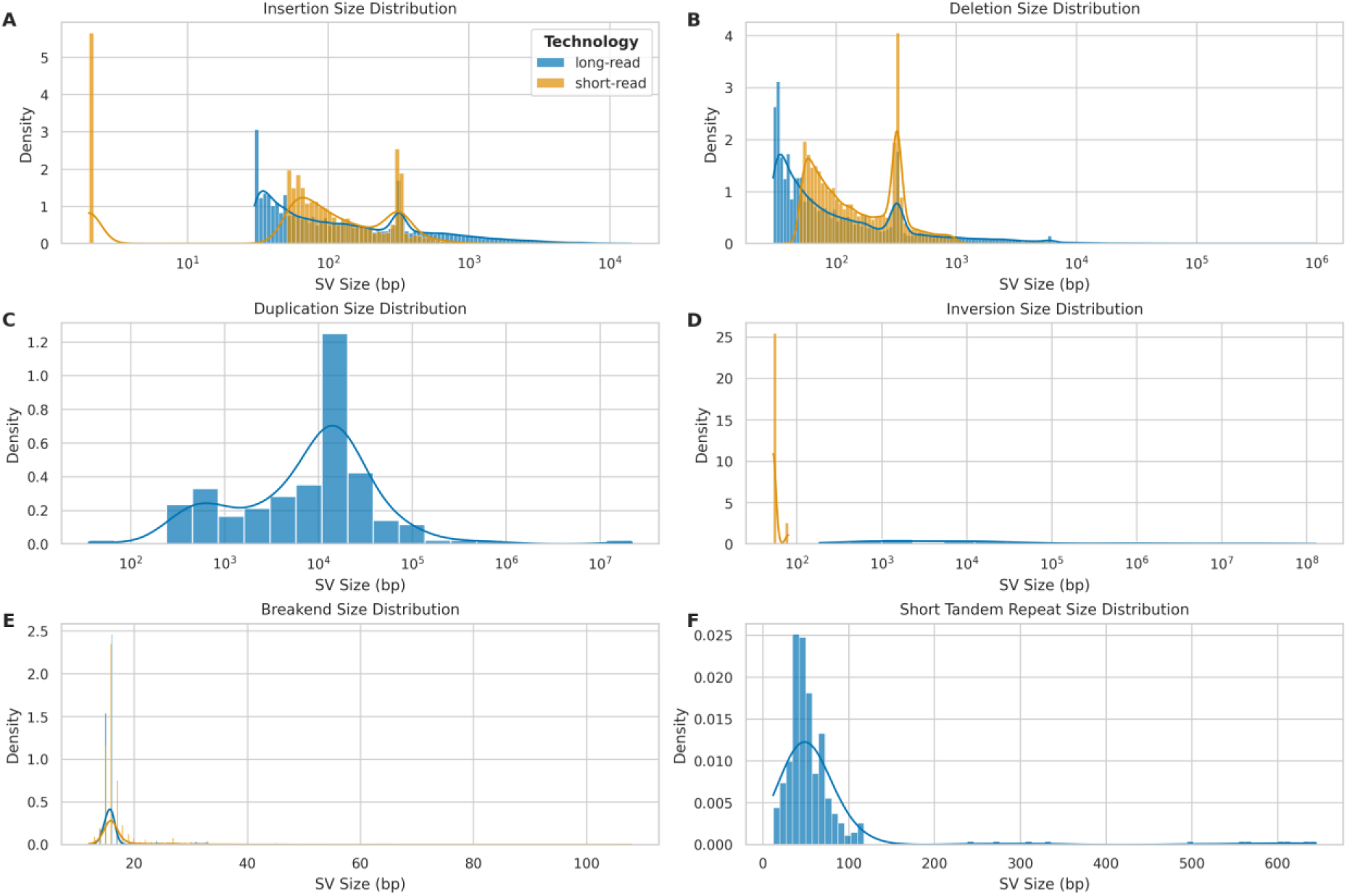
Size distribution of different structural variant (SV) types detected by long-read (ONT) and short-read (Illumina) sequencing platforms. **(A)** Insertion size distribution: long-read (blue) vs. short-read (orange). **(B)** Deletion size distribution: long-read (blue) vs. short-read (orange). **(C)** Duplication size distribution: primarily detected by long-read platform (blue). **(D)** Inversion size distribution: long-read (blue) vs. short-read (orange). **(E)** Breakend (BND) size distribution: long-read (blue) vs. short-read (orange). **(F)** Short Tandem Repeat (STR) size distribution: exclusively detected by long-read platform (blue).

Duplication events (**Figure 12C**) were primarily detected by LRS, with sizes ranging from 100 bp to 100 Mb and a peak around 10 kb. The absence of a substantial SRS signal for duplications highlights the advantage of LRS in identifying these complex structural variants. Inversion size distributions (**Figure 12D**) showed striking differences between the platforms. LRS detected inversions across a broad size range (10² to 10⁸ bp), while SRS mainly identified smaller inversions (< 10³ bp).

Breakend (BND) size distributions (**Figure 12E**) were similar between platforms, with both showing a peak around 20 bp. However, SRS demonstrated a slightly broader range, extending to approximately 100 bp. Short Tandem Repeat (STR) size distributions (**Figure 12F**) were exclusively captured by LRS, with sizes ranging up to approximately 700 bp and a peak around 50 bp.

To investigate the genomic distribution of SVs, we analysed their occurrence across chromosomes, normalizing for chromosome length (**Figure 13A**). For both LRS and SRS platforms, chromosomes 19, 20, 21 exhibited notably higher percentages of SVs per megabase compared to other chromosomes, while, conversely, chromosomes 14, 15, X, and Y showed lower SV densities for both platforms. Interestingly, despite their technological differences, both LRS and SRS platforms showed similar trends in chromosomal SV distribution. A further correlation analysis between chromosome length and SV occurrence (**Figure 13B**) revealed weak and non-significant correlations between chromosome length and SV counts for both LRS (r = 0.11, p = 0.616) and SRS (r = 0.05, p = 0.824) platforms, suggesting that chromosome length is not a strong predictor of SV abundance.

**Figure 13.**
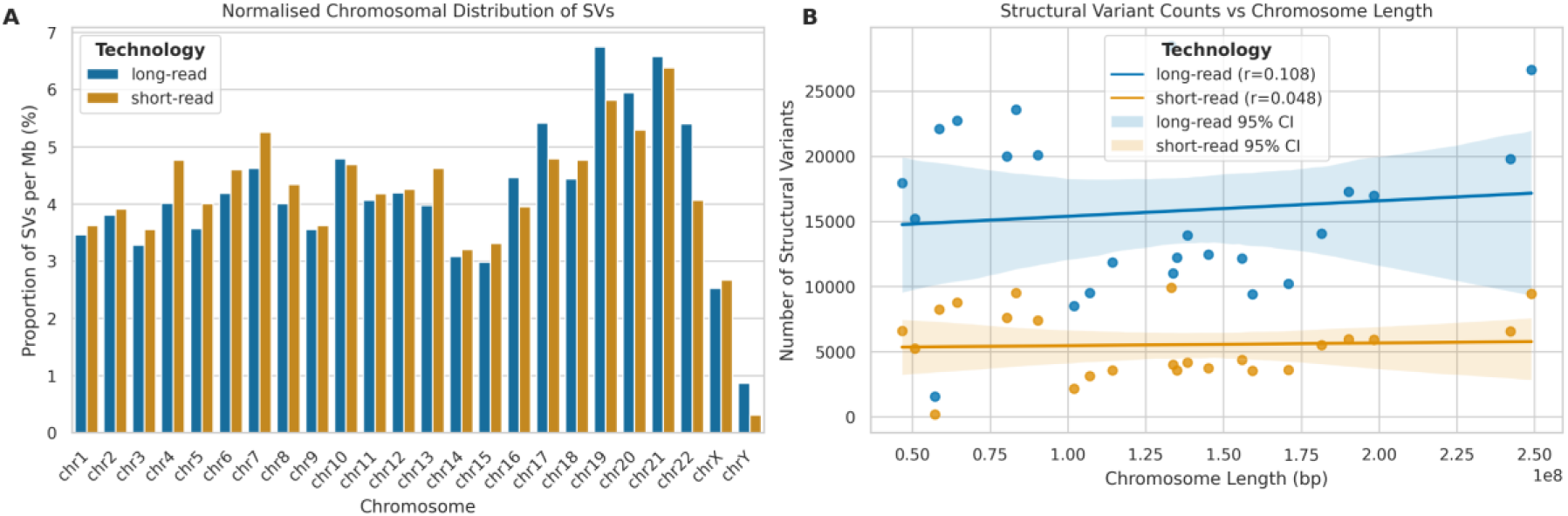
Chromosomal distribution and correlation analysis of SVs detected by long-read (ONT) and short-read (Illumina) sequencing. **(A)** Normalised chromosomal distribution of SVs (% per Mb) for long-read (blue) and short-read (orange) platforms. **(B)** Correlation between SV counts and chromosome length (bp) for both platforms. Lines represent best fit with 95% confidence intervals. Pearson correlation coefficients (r) provided.

The elevated SV density observed in chromosomes 19, 20, and 21 could be attributed to various factors, including higher gene density, increased recombination rates, or the presence of specific genomic features that promote structural rearrangements (41). For instance, chromosome 19, is known for its high gene density and elevated GC content, which may contribute to its increased propensity for structural variations (42).

### Impact of sequencing depth on calling structural variants

To evaluate the influence of sequencing depth on structural variant (SV) detection, we analysed the relationship between whole genome mean depth and both the number and size distribution of SV calls across LRS and SRS platforms.

For LRS, we observed a strong positive correlation between sequencing depth and the number of SV calls (**Figure 14A**) (r = 0.94, p = 6.71×10^-7^). The relationship followed an asymptotic trend, with the number of SV calls increasing from approximately 16,000 at 10X coverage to around 35,000 at 30X coverage, though the rate of increase diminished at higher depths. The consensus calls between the two platforms demonstrated a strong positive correlation with sequencing depth (**Figure 14B**) (r = 0.89, p = 2.14×10^-5^), suggesting that increased depth primarily improves the detection of true structural variants. Analysis of SV size distributions across sequencing depths revealed that for LRS, we observed consistent detection of SVs across a broad size range (10¹ to 10⁸ bp) at all sequencing depths, with no significant trend in median SV size (r = 0.006, p = 2.17×10-4). The mean SV size showed considerable variation (range: 412 - 9,848 bp), likely due to improved detection of larger variants at higher depths (**Figure 14C**).

**Figure 14.**
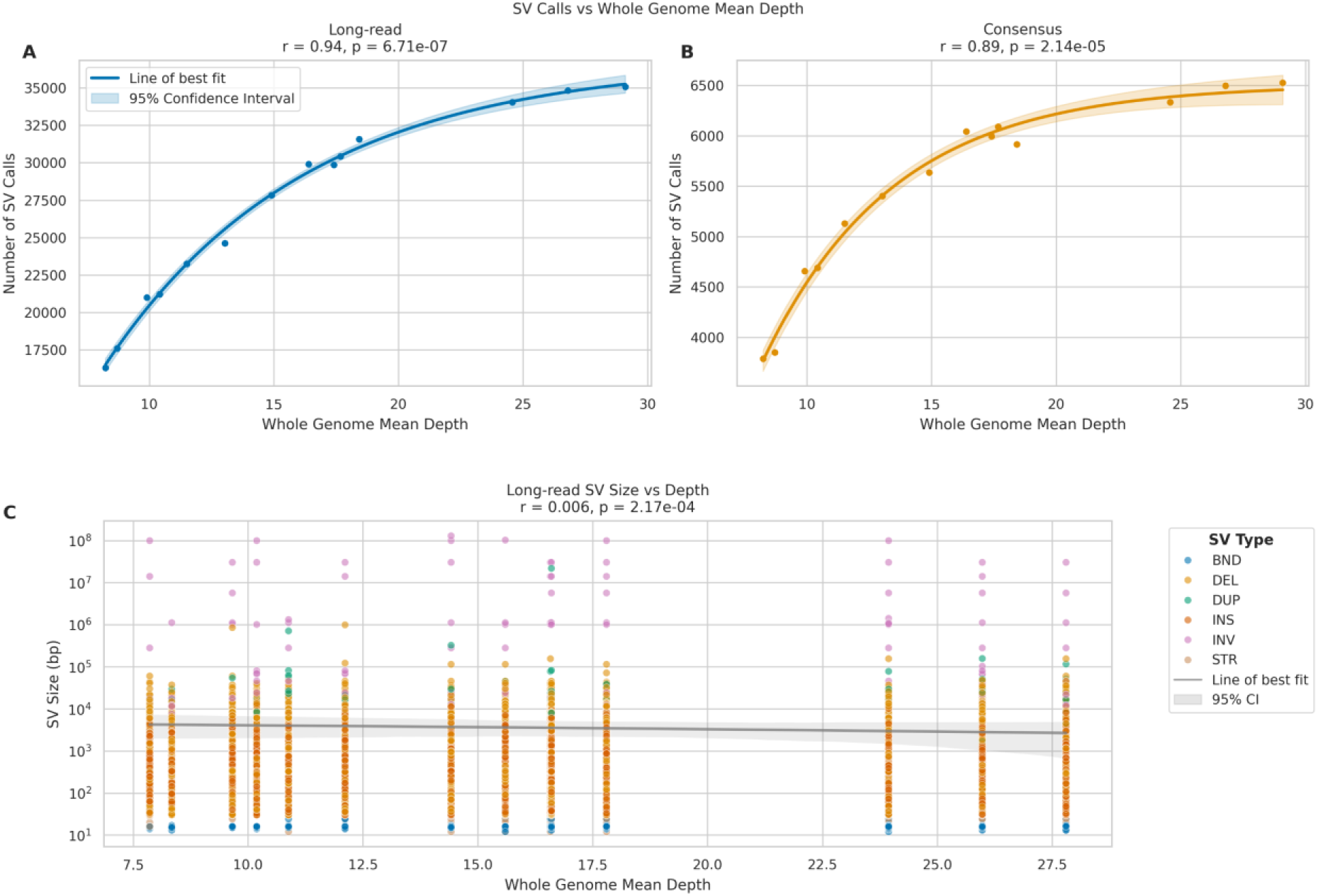
Impact of sequencing depth on SV detection in long-read (ONT). **(A, B)** Scatter plots relating whole genome mean depth to SV call numbers for long-read (A) and consensus calls (B). Lines show best fit with 95% confidence intervals (shaded). Pearson correlation coefficients (r) and p-values provided. **(C)** SV size distribution across sequencing depths for long-read data. Y-axis shows SV size (log scale). Colours indicate SV types: INS (insertions), DEL (deletions), DUP (duplications), INV (inversions), BND (breakends), and STR (short tandem repeats).

These findings suggest that while increased sequencing depth substantially improves SV detection in long-read sequencing, particularly for larger variants, short-read SV calling remains relatively consistent across different depths. The data indicate that a minimum of 25X coverage may be optimal for long-read SV detection, as the rate of novel SV discovery plateaus beyond this depth.

## Discussion

This study provides a comprehensive comparison of Oxford Nanopore Technologies (ONT) long-read sequencing (LRS) and Illumina short-read sequencing (SRS) platforms, focusing on their performance in variant detection across different genomic contexts. Our findings provide a comprehensive understanding of their performance in variant detection, particularly in challenging genomic regions, and highlight the impact of experimental factors on sequencing outcomes.

### Sequencing Quality and Yield

The comparison between multiplexed and singleplexed ONT samples revealed significant differences in sequencing yield and read length characteristics. Singleplexed samples consistently produced higher numbers of reads and bases, with a 118.48% increase in mean read count and a 99.94% increase in mean base count compared to multiplexed samples. This substantial difference in yield has important implications for experimental design and cost-effectiveness in genomic studies. While multiplexing offers cost savings and increased throughput, our results suggest that it may come at the expense of reduced per-flowcell sequencing depth and potentially slightly lower variant detection accuracy, particularly for structural variants and in low complexity regions, although this can be mitigated by optimising barcoding.

Interestingly, we observed distinct read length distribution patterns between multiplexed and singleplexed samples. The consistent bimodal distribution in singleplexed samples, with peaks at shorter (∼400 bp) and longer (∼10,000 bp) read lengths, contrasts with the more variable patterns seen in multiplexed samples. This difference may reflect variations in library preparation or sequencing efficiency between the two approaches and warrants further investigations to optimise multiplexing protocols for long-read sequencing.

### Single Nucleotide Variants Detection

Our analysis of SNV detection performance revealed that while SRS maintains an overall advantage, particularly in high complexity regions, LRS demonstrates competitive performance, especially in low complexity areas. In high complexity regions, SRS showed statistically significant higher precision, sensitivity, and F-measure compared to LRS. However, the performance gap narrowed considerably in low complexity regions, with LRS showing comparable sensitivity to SRS.

The distinct error profiles observed for LRS and SRS across different SNV types provide valuable insights into the strengths and weaknesses of each technology. The higher error rates for LRS, particularly for transitions in high complexity regions, suggest that improvements in base-calling accuracy should remain a priority for ONT. However, the competitive performance of LRS in detecting certain SNV types within low complexity regions, particularly in terms of false negative rates, highlights its potential advantages in analysing repetitive genomic areas where traditional short-read sequencing faces inherent limitations (22).

### Indels Detection

The performance of LRS in indel detection showed marked differences between high and low complexity regions, with a clear dependency on indel size. In high complexity areas, LRS demonstrated robust performance particularly for small (1-5bp) and medium-sized indels (6-20bp), which represent the vast majority of variants (>95%), achieving F-measures of up to 0.869. However, there was a significant decline in performance in low complexity regions, where even small indels showed markedly reduced F-measures of 0.454. Performance deteriorated further with increasing indel size in these regions, with F-measures dropping to 0.350 for indels of 21-50bp.

This size-dependent pattern suggests that while current LRS technology can reliably detect small to medium-sized indels in non-repetitive regions, significant challenges remain in accurately calling indels of any size in repetitive areas. The particularly poor performance for larger indels in low complexity regions could partially reflect limitations in our SRS-based benchmarking approach, as short-read technologies face inherent challenges in detecting larger variants in repetitive regions. Future benchmarking studies incorporating more robust validation methods may help distinguish true technological limitations from artifacts of the comparison methodology.

### Impact of Experimental Factors on Variant Calling

Our investigation into the effects of multiplexing, sequencing depth, and read length on variant calling performance yielded several important insights. Initially, multiplexing appeared to have a modest impact on SNV calling performance and a more substantial effect on indel calling accuracy. However, our analysis revealed that these apparent effects were largely attributable to differences in sequencing depth rather than multiplexing status itself. After controlling for depth, multiplexing showed minimal impact on SNV calling and only modestly affected indel precision. This finding suggests that researchers can confidently employ multiplexing strategies without significantly compromising variant calling accuracy, provided adequate sequencing depth is maintained. The ANCOVA results demonstrated particularly strong relationships between sequencing depth and variant calling performance metrics, especially for indel detection (r² > 0.95). This finding highlights the importance of selecting adequate coverage in genomic studies. Furthermore, the asymptotic trend in this relationship suggests that while increasing sequencing depth generally improves variant calling performance, there may be diminishing returns at very high depths. This information can guide researchers in optimising their sequencing strategies to balance cost and variant calling accuracy.

Notably, we also found a moderate negative correlation between mean read length and variant calling performance for both SNVs and indels. This unexpected result warrants further investigation and may reflect interactions between read length, sequencing errors, and alignment challenges. It also underscores the need for continued refinement of bioinformatics tools to fully leverage the potential of the read lengths generated by LRS.

### Structural Variants Detection

Our analysis revealed significant differences in SV detection capabilities between LRS and SRS platforms. LRS identified 2.86 times more SVs compared to SRS across all samples, demonstrating its enhanced ability to capture a broader spectrum of structural variations. This advantage was particularly evident at higher sequencing depths, with our analysis revealing a strong positive correlation between depth and SV detection in LRS (r = 0.94, p = 6.71×10^-7^). The relationship in LRS followed an asymptotic trend, suggesting optimal SV detection might be achieved around 25X coverage. Furthermore, the superior capability of LRS in detecting large-scale genomic rearrangements, as evidenced by the markedly larger maximum SV size (129 Mb for LRS vs 6 Kb for SRS), highlights its potential in uncovering previously undetectable genomic events. Importantly, this broad size range detection capability remained consistent across all sequencing depths in LRS (10 to 10⁸ bp), while SRS maintained a more constrained size distribution (10 to 1000 bp) regardless of coverage depth.

The distinct patterns observed in SV size distributions and types between LRS and SRS platforms further emphasise their complementary nature. While SRS showed a bias towards detecting smaller SVs, LRS demonstrated a broader range of detectable SV sizes and types, including improved detection of large insertions, deletions, and inversions. The exclusive detection of Short Tandem Repeats (STRs) by LRS underscores its unique advantages in detecting certain types of genomic variation. However, it is important to note that these differences are not solely attributable to the sequencing technologies themselves, but also to the specific variant callers used for each platform in this study. Indeed, the choice of variant caller can significantly impact the number and types of structural variants discovered, and utilising different callers or a combination of multiple callers on the same data could potentially increase the number and diversity of structural variants identified on both platforms. Future studies comparing LRS and SRS technologies should consider evaluating and comparing the performance of multiple variant callers to provide a more comprehensive assessment of each platform’s capabilities in structural variant detection.

In addition, the analysis of chromosomal distribution of SVs revealed interesting patterns, with chromosomes 19, 20, and 21 exhibiting notably higher SV densities, and the strong positive correlation between chromosome length and SV counts for both platforms provides further insights into the genomic landscape of structural variations.

### Limitations and Future Directions

While our study provides insights into the performance of ONT LRS technology, several limitations should be acknowledged. Our deliberate selection of the highest quality flowcells for multiplexing introduced a methodological bias that confounds the direct comparison between multiplexed and singleplexed approaches. This bias, while potentially minimising the performance impact of multiplexing, makes it challenging to fully assess the true effects of multiplexing on sequencing quality and variant detection. Moreover, our relatively small sample size (n=14) limits the statistical power and generalisability of our findings, and the use of SRS as a gold standard for indel comparisons may introduce biases, particularly in regions where SRS itself faces challenges. Additionally, the potential biases in SV detection methods and the limited sample size and diversity in our study call for further investigations with larger, more diverse cohorts.

To address these limitations, future studies should employ randomized flowcell allocation for multiplexing experiments and significantly larger sample sizes to provide more robust statistical comparisons. A larger comprehensive study, using randomly assigned flowcells of varying quality, would better elucidate the true impact of multiplexing and flowcell quality on sequencing performance. Such a study could also incorporate technical replicates to assess the reproducibility of findings across different flowcell qualities.

Looking ahead, the integration of LRS and SRS technologies presents exciting opportunities for comprehensive genomic analysis. The complementary strengths of these platforms could enable more accurate and complete characterisation of genomic variations across different scales and complexities. Continued improvements in base-calling accuracy and variant detection algorithms for LRS will be vital in fully realising its potential. Future research should explore the applications of LRS in clinical genomics and complex trait studies, where its ability to detect a wider range of structural variations could uncover novel disease-associated variants. Additionally, further optimisation of multiplexing protocols and sequencing strategies for LRS could enhance its cost-effectiveness and accessibility for large-scale genomic studies.

## Methods

### Sample Selection and Preparation

Samples used in this study were obtained from Project MinE, an international collaboration focused on whole-genome sequencing of amyotrophic lateral sclerosis (ALS) patients and controls (43). From this cohort, we selected 14 samples for analysis. Venous blood was drawn from participants, and genomic DNA was isolated using standard methods. DNA concentrations were standardized to 100 ng/μl as measured by fluorometry using the Quant-iT™ PicoGreen™ dsDNA quantitation assay (Thermo Fisher Scientific). DNA integrity was assessed using gel electrophoresis.

### Microarray Genotyping

DNA samples were genotyped using the Infinium Omni2.5-8 v1.4 array (Illumina, Cambridge, United Kingdom) according to the manufacturer’s protocol. To ensure consistency across platforms, the genotyped variants were lifted over to the GRCh38 reference genome using GATK LiftoverVcf (44). Only variants that were successfully lifted over and had matching alleles in the reference genome were retained for subsequent analyses.

### Illumina Short-Read Sequencing

DNA samples were sequenced using Illumina’s FastTrack services (Illumina, San Diego, CA, USA) on the Illumina HiSeq 2000 platform. Sequencing was performed using 100 bp paired-end reads with PCR-free library preparation, yielding approximately 35× coverage across each sample. The Isaac pipeline (45) was used for alignment to the hg19 reference genome and to call single nucleotide variants (SNVs), insertions and deletions (indels), and larger structural variants (SVs). To ensure consistency across platforms, the called variants were lifted over to the GRCh38 reference genome using GATK LiftoverVcf (44). Only variants that were successfully lifted over and had matching alleles in the reference genome were retained for subsequent analyses.

### Oxford Nanopore Long-Read Sequencing

For ONT sequencing, DNA libraries were extracted with the Monarch® Spin gDNA Extraction Kit T3010S (New England Biolabs) and prepared using the SQK-LSK114 Ligation Sequencing Kit (Oxford Nanopore Technologies), according to the respective manufacturers’ protocols. The prepared libraries were quantified using a Qubit fluorometer (Thermo Fisher Scientific). For multiplexed runs, barcoding was performed using the SQK-NBD114.24 Native Barcoding Kit (Oxford Nanopore Technologies). Sequencing was performed on PromethION 24 instruments using R10.4.1 flowcells. Two samples were pooled and sequenced on a single flow cell for multiplexed runs, while individual samples were sequenced on dedicated flow cells for singleplexed runs. Sequencing runs were performed for approximately 72 hours.

Raw signal data were base-called with the EPI2ME wf-basecalling v1.4.5 pipeline (46), using the Dorado model dna r10.4.1 e8.2 400bps sup v5.0.0. The unaligned BAM output was assessed for quality with NanoPlot v1.42.0 (47). Using the EPI2ME wf-human-variation v2.6.0 pipeline (48-56), reads were aligned to the GRCh38 reference genome; this pipeline uses Clair3 (51) to call SNVs and indels, Sniffles2 (52) to detect SVs, and Straglr (56) for STRs.

### Variant Comparison and Benchmarking

All variant files were filtered and compared using a custom Nextflow pipeline, which is available at https://github.com/renatosantos98/ont-benchmark and can be executed using the command *nextflow run renatosantos98/ont-benchmark*.

### Single Nucleotide Variants and Indels

For SNVs, variants from all three platforms (ONT LRS, Illumina SRS, and Illumina microarray) were intersected with regions excluding and including the Tandem Repeats and Homopolymers regions defined by the GIAB v3.4 benchmark (33), to form the high complexity and low complexity regions, respectively. Only variants present in the microarray dataset were retained for the LRS and SRS datasets to ensure consistent comparison across platforms.

For indels, variants from LRS and SRS platforms were also filtered, based on the same high-confidence regions, into high-complexity and low-complexity sets based on their genomic location.

Comparison of SNVs and indels was performed using RTG Tools vcfeval v3.12 (37, 38), to generate true positive, false positive, and false negative variant sets, as well as performance metrics including precision, recall, and F-measure. For SNV comparisons, the microarray data served as the truth set, while for indel comparisons, the Illumina SRS data was used as the truth set.

### Structural Variants

SVs from LRS and SRS were filtered for quality, whereby for LRS variants with at least 5 supporting reads and STRs with a minimum of 10 spanning reads were retained, and for SRS variants with at least 5 supporting paired-end or split reads were retained.

The filtered callsets were compared using SURVIVOR v1.0.7 (40), by generating a merged consensus set of variants from both platforms which included only variants within 500 base pairs of each other, of the same type, on the same strand, and with a size difference of no more than 30%.

### Statistical Analysis

Statistical analyses were performed using jupyterlab 4.3.5 (57), Python 3.12.8 (58), numpy 2.2.2 (59), polars 1.21.0, matplotlib 3.10.0 (60), seaborn 0.13.2 (61), SciPy 1.15.1 (62), scikit-learn 1.6.1 (63), and statsmodels 0.14.4 (64). This work was performed on the King’s Computational Research, Engineering and Technology Environment (CREATE) at King’s College London (65). Statistical significance was deemed as a p-value < 0.05.

## Supplementary Materials

The source code used to generate the results presented in this study is available in the GitHub repository https://github.com/renatosantos98/ont-benchmark.

## Informed Consent Statement

Informed consent was obtained from all subjects involved in the study.

## Data Availability

The raw data supporting the conclusions of this article will be made available by the authors on request.

